# Tissue and cell-type specific expression of a splice variant in the II-III cytoplasmic loop of *Cacna1b*

**DOI:** 10.1101/596668

**Authors:** Bunda Alexandra, LaCarubba Brianna, Akiki Marie, Andrade Arturo

**Author notes:** Corresponding author: 46 College Road, 214 Rudman Hall. Durham, NH, USA. 03824. Both authors contributed equally to this work (Bunda Alexandra, LaCarubba Brianna).

## Abstract

Presynaptic Ca_V_2.2 (N-type) channels are fundamental for transmitter release across the nervous system. The gene encoding Ca_V_2.2 channels, *Cacna1b*, contains alternatively spliced exons that originate functionally distinct splice variants (e18a, e24a, e31a and 37a/37b). Alternative splicing of the cassette exon 18a generates two mRNA transcripts (+e18a-*Cacna1b* and Δe18a-*Cacna1b*). In this study, using novel mouse genetic models and in situ hybridization (BaseScope™), we confirmed that +e18a-*Cacna1b* splice variants are expressed in monoaminergic regions of midbrain. We expanded these studies and identified +e18a-*Cacna1b* mRNA in deep cerebellar cells and spinal cord motor neurons. Furthermore, we determined that +e18a*-Cacna1b* is enriched in cholecystokinin expressing interneurons. Our results provide key information to understand cell-specific functions of Ca_V_2.2 channels.

## INTRODUCTION

Presynaptic Ca_V_2.2 (N-type) channels are key mediators of transmitter release across the nervous system. The calcium that enters through Ca_V_2.2 channels in response to action potentials triggers transmitter release [1, 2]. *Cacna1b* is a multi-exon gene that encodes the Ca_V_*α*_1_ pore-forming subunit of Ca_V_2.2 channels. The *Cacna1b* pre-mRNA contains more than 40 exons that are spliced during mRNA maturation [3]. Most *Cacna1b* exons are constitutively included in the final *Cacna1b* mRNA, however some are selectively included through alternative splicing (e18a, e24a, e31a and 37a/37b) [4]. Alternative splicing of the *Cacna1b* pre-mRNA originates splice variants that are translated into Ca_V_2.2 channels with distinct biophysical properties and pharmacological profiles [5, 6]. Ca_V_2.2 splice variants with differences in activation, inactivation and current density [7–9], as well as in their response to neurotransmitters and drugs such as GABA and morphine have been previously characterized [10, 11]. Thus, alternative splicing of the *Cacna1b* pre-mRNA is thought to provide functional diversification to cells that utilize Ca_V_2.2 channels to release neurotransmitter.

Among the alternative spliced exons in the *Cacna1b* pre-mRNA is the cassette exon 18a (e18a) [12]. Alternative splicing of e18a generates two splice variants, +e18a-*Cacna1b* (e18a is included) and Δe18a*-Cacna1b* (e18a is skipped) (**Fig. 1A**). E18a encodes 21 amino acids within the “synprint” region of the II-III cytoplasmic loop of Ca_V_2.2 (LII-III), an essential region for interaction of Ca_V_2.2 channels with presynaptic proteins [13–15]. Functionally, +e18a-Ca_V_2.2 channels are more resistant to cumulative inactivation induced by repetitive stimulation and closed-state inactivation than Δe18a-Ca_V_2.2 channels [9]. Furthermore, +e18a-Ca_V_2.2 channels exhibit larger calcium currents relative to Δe18a-Ca_V_2.2 channels in both mammalian expression systems and neurons [16]. Despite all of this information, very little is known about the functional role of +18a-*Cacna1b* and Δe18a-*Cacna1b* splice variants in the nervous system.

**Figure 1.**
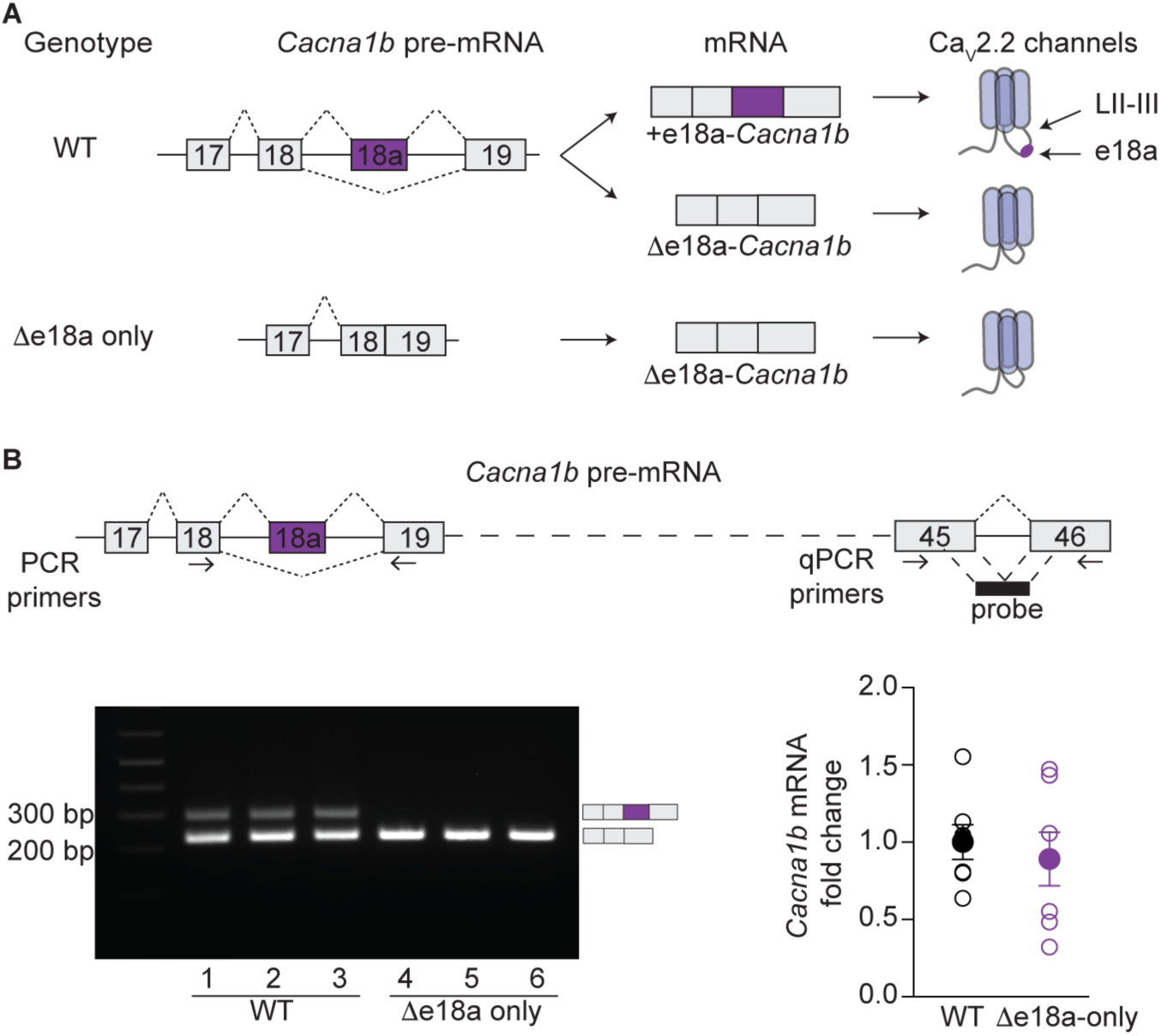
Targeted deletion of e18a in the Cacna1b gene. **A**) Schematic of splicing patterns for *Cacna1b* (Ca_V_2.2) pre-mRNA in WT and Δe18a-only mice. In WT mice, e18a is spliced to generate two mRNA transcripts, +e18a- and Δe18a-*Cacna1b*, which in turn are translated into two different Ca_V_2.2 channels, +e18a- and Δe18a-Ca_V_2.2. The *Cacna1b* gene was modified to remove e18a and its flanking intronic regions to generate eΔ18a-only mice, which generate only the Δe18a-Ca_V_2.2 splice variant. **B**) *Top*, Schematic of *Cacna1b* pre-mRNA. Arrows indicate the approximate location of RT-PCR primers flanking e18a and qPCR primers in constitutive exons 45 and 46. Black box shows the approximate location of qPCR probe spanning exon junction 45-46. *Bottom left*, RT-PCR from whole brain samples of WT and Δe18a-only mice. *Bottom right*, comparison of whole brain *Cacna1b* mRNA levels between WT and Δe18a-only mice. Data are shown as mean (filled symbols) ± s.e.m, and individual values for biological replicates (empty symbols).

To understand the functional role of +e18a-*Cacna1b* and Δe18a-*Cacna1b* splice variants, it is necessary to determine their tissue and cell-type expression. Previous studies have shed light on this. In the adult nervous system, the abundance of +e18a-*Cacna1b* mRNA differs among tissues. Higher levels of +e18a-*Cacna1b* mRNA are observed in dorsal root ganglia, superior cervical ganglia, and spinal cord relative to whole brain [6, 16]. Interestingly, the relative abundance between +e18a-*Cacna1b* and Δe18a-*Cacna1b* mRNAs also differs among brain subregions. Less than 10% of *Cacna1b* splice variants contain e18a in whole cerebral cortex and hippocampus, whereas 30-60% of *Cacna1b* splice variants contain e18a in the thalamus, cerebellum, hypothalamus and midbrain [12]. Within the midbrain, ∼80% of *Cacna1b* splice variants contain e18a in monoaminergic regions including the ventral tegmental area (VTA), substantia nigra (SN), dorsal raphe nuclei (DRN), and locus coeruleus (LC) [17].

The expression of +e18a-*Cacna1b* is also cell-specific. +e18a-*Cacna1b* mRNA is enriched in tyrosine hydroxylase-expressing cells of the SN pars compacta (SNc) and VTA [17]. +e18a-*Cacna1b* has also been identified in magnocellular neurosecretory cells of hypothalamus and capsaicin-responsive neurons of dorsal root ganglia [7, 18]. The molecular mechanisms underlying the tissue- and cell-specific expression of +e18a-*Cacna1b* and Δe18a-*Cacna1b* mRNAs are beginning to be elucidated. The RNA-binding protein, Rbfox2, is a splicing factor that represses e18a by binding an intronic region upstream of this exon [16, 19]. Rbfox2 expression and activity depend on the cell-type; thus, this would help to explain the cell-specific expression of +e18a-*Cacna1b* and Δe18a-*Cacna1b* [20, 21].

In this study, we utilized a novel version of in situ hybridization (BaseScope™) and genetic mouse models, to precisely determine the expression of +e18a-*Cacna1b* in the central nervous system. We generated a mouse model that lacks expression of +e18a-*Cacna1b*, thereby expresses only the Δe18a-*Cacna1b* splice variant (Δe18a-only). This mouse was used to control for probe specificity in our BaseScope™ experiments. With these new approaches and tools, we confirmed that +e18a-*Cacna1b* mRNA is expressed in monoaminergic regions (SN, VTA, DRN, and LC). Furthermore, we identified new cell populations that express +e18a-*Cacna1b* mRNA such as cells of the deep cerebellar nuclei (DCN) and spinal cord motor neurons. Finally, using fluorescence-activated cell sorting (FACS) of genetically identified cell populations coupled to RT-PCR, we found that +e18a-*Cacna1b* mRNA is more abundant in cholecystokinin expressing interneurons (CCK+INs) relative to Ca2+/calmodulin-dependent protein kinase II*α* expressing projection neurons (CaMKII*α*+PNs).

## MATERIALS AND METHODS

### Housing Conditions

A combination of adult males or females were used in all of our experiments, no association to sex was found in the amount of e18a in *Cacna1b* in whole brain. Mice were housed with food and water *ad libitum* in temperature-controlled rooms with a 12 h light/dark cycle. All experimental procedures followed the guidelines of the Institutional Animal Care Committee of the University of New Hampshire.

### Mouse lines

C57BL/6 wild-type mice were used in our experiments. Mice lacking e18a or Δe18a-only (*Cacna1b^tm4.1Dili^*) were back crossed in C57BL/6 (Charles River) background for 6-8 generations, for details on how this mouse line was generated see [16]. To label CCK+INs with tdTomato (tdT), we performed intersectional genetic labeling as previously reported [22, 23]. We utilized *CCK-Cre* (*Cck*^*tm1.1(cre)Zjh*^/J, Jax:012706) in C57BL/6 background, *Dlx5/6-Flpe* (*Tg(mI56i-flpe)39Fsh/J*, Jax: 010815) in FVB/NJ background, and *Ai65-D* (*B6;129S-Gt(ROSA)26Sort^m65.1(CAG-tdTomato)Hze^/J*, Jax: 021875) in C57BL/6;I129 background mice. Using these three mouse lines, we generated a triple transgenic line, *CCK-Cre; Dlx5/6-Flpe; Ai65-D* (named *CCK;Dlx5/6;tdT*) as follows: First, *CCK-Cre* mice were crossed with *Dlx5/6-Flpe* two times to produce a dual transgenic mouse (homozygous for *CCK-Cre* and heterozygous for *Dlx5/6-Flpe*). Next, this dual transgenic mouse line was crossed with homozygous *Ai65-D* mice. From the resulting offspring, we selected only heterozygous mice for the three alleles. To induce the expression of tdT in PNs, we crossed *CaMKIIα-Cre* mice (*B6.Cg-Tg(Camk2a-cre)T29-1Stl/J*, Jax: 005359) in a mixed C57BL6 background with *Ai14* mice (*B6.Cg-Gt(ROSA)26Sor^tm14(CAG-tdTomato)Hze^/J*, Jax: 007914) in C57BL6. The resulting dual transgenic mouse line *CaMKIIα;tdT* was heterozygous for both alleles.

### Genotyping

Conventional toe biopsy was performed on P7-P9 pups. Genomic DNA was extracted using Phire Animal Tissue Direct kit II (ThermoFisher Scientific, F140WH) according to manufacturer instructions. Next, PCR was performed with AmpliTaq Gold^®^ 360 mastermix (Thermo Fisher Scientific) using the following conditions: a hot start of 95°C for 10 min, followed by 35 cycles (95°C, 30 s; 60°C, 30 s; 72°C, 1 min), and final step of 72°C for 7 min. Primers and expected products are shown in Table 1. Primers were added together to genotype each mouse line.

### In situ hybridization (BaseScope™)

Mice were deeply anesthetized with Euthasol (Virbac, 200-071). Next, mice were transcardially perfused with 1x PBS for 10 min and subsequently with 10% neutral buffered formalin (NBF, ∼4% formaldehyde) fixative solution (Sigma, HT501128) for 10 min, and then again with 1x PBS for 10 min. Both PBS and NBF were kept on ice during the perfusion. Brain and spinal cord were immediately removed and post-fixated in 10% NBF at 4°C for 24 hours. After washing with 1x PBS, tissue was sequentially dehydrated in 15% and 30% sucrose:PBS solutions for at least 18 hours in each sucrose concentration, or until tissue sank to the bottom of the tube. Next, tissue was cryopreserved in optimal cutting temperature compound, or OCT (Fisher, 4585), with isopentane pre-chilled in dry ice. 12 μm cryosections from brain and spinal cord were collected (Shandon, 77200222) and placed in pre-chilled 15 mm Netwell™ inserts (Corning, 3478). Sections were allowed to free-float in 1x PBS, and mounted on positively charged microscope slides (VWR, 48311-703) using a paintbrush. After air-drying for 10-20 min, sections were incubated at 60°C in a drying oven for 30 minutes to aid tissue adhesion to the slides. Before *in-situ* hybridization (ISH), sections were post-fixed in 10% NBF at 4° C for 15 minutes, then dehydrated with 50%, 70%, and two rounds of 100% ethanol for 5 minutes in sequential steps. Slides were air-dried for an additional 5 minutes before incubating with RNAscope® hydrogen peroxide solution for 10 minutes (ACD, 322381). Next, sections were washed with milliQ water and transferred to RNAscope® Target Retrieval solution preheated to 99° C for 15 minutes (ACD, 322000). Sections were briefly washed with milliQ water, transferred to 100% ethanol for 3 minutes, then placed in drying oven at 60°C for 30 minutes. Sections were isolated with a hydrophobic barrier pen (ACD, 310018) and air-dried overnight at room temperature. Following incubation with RNAscope® Protease III for 30 minutes (ACD, 322381) at 40° C in the ACD HybEZ™ Hybridization System (ACD, 310010), sections were exposed to a probe spanning e18 and e18a (BaseScope™ Mm-Cacna1b-e18e18a) (ACD, 701151) for 2 hours at 40° C in the ACD HybEZ™ Hybridization System. BaseScope™ Detection Reagents AMP 0 - AMP 6 and FastRed (ACD, 322910) were applied according to manufacturer instructions and washed using RNAscope® Wash Buffer (ACD, 310091). To visualize nuclei, sections were counterstained in Gill’s Hematoxylin I for 2 minutes at RT (American Master Tech, HXGHE). Next, sections were washed with tap water 3 times, briefly transferred to 0.02% ammonia water, and washed again with tap water. Finally, sections were dried at 60° C for 15 minutes and mounted using VectaMount™ mounting medium (Vector Laboratories, H-5000). Phase contrast images were acquired using an Olympus IX-81 microscope.

### Fluorescence-activated cell sorting

Adult *CCK;Dlx5/6;tdT* and *CaMKIα;tdT* mice, were deeply anesthetized with isofluorane. After decapitation, brains were quickly removed and placed on a petri dish with Earl’s Balanced Salt Solution (EBSS) (Sigma, E3024) containing 21 U/mL of papain. Rapid dissection (< 45 s) of cerebral cortex and hippocampus was performed. Then, tissue was dissociated using a modified version of Worthington Papain Dissociating System® (Worthington Biochemical Corporation, LK003150). After incubating with papain for 45 min at 37°C on a rocking platform, tissue was triturated with three sequential diameter fire-polished glass pipettes. Next, cell suspensions were centrifuged at 300 g for 5 min. After discarding supernatants, pellets were resuspended in 3 mL of EBSS containing 0.1 % of ovomucoid protease inhibitor and 0.1 % bovine serum albumin (Worthington, LK003182) to quench papain. Cell suspension was centrifuged at 270 g for 6 min and resuspended in EBSS (3 mL). To isolate tdT-expressing cells, we performed FACS in a Sony SH800 flow cytometer using a 561 nm laser to excite and a 570-630 nm filter for event selection. At least 300,000 events were collected directly into TRIzol™ LS Reagent (ThermoFisher Scientific, 10296028). Collection was performed keeping 1:3 (v/v) sorted cell suspension: TRIzol™ LS ratio. Cell suspension was kept on ice throughout the sorting session.

### RT-PCR and RT-qPCR

Total RNA from tissue was extracted with RNeasy Mini Kit columns (Qiagen, 74134) according to manufacturer instructions. Total RNA from sorted cells was extracted using TRIzol LS and isopropanol precipitation with the addition of 30 μg of GlycoBlue® Coprecipitant (ThermoFisher Scientific, AM9516) to facilitate visualization of RNA pellet. 1 μg (tissue) or 300 ng (sorted cells) of total RNA was primed with oligo-dT and reverse transcribed with Superscript IV First-Strand Synthesis System (ThermoFisher Scientific, 18091050) according to manufacturer instructions. To quantify the relative amount of e18a, PCR was performed using AmpliTaq Gold^®^ 360 master mix (ThermoFisher Scientific, 4398881) with primers flanking e18a (F: 5’GGCCATTGCTGTGGACAACCTT and R: 5’CGCAGGTTCTGGAGCCTTAGCT) with the following conditions: Hot start at 95°C for 10 min, 28 cycles (95°C for 30 sec, 60°C for 30 sec, 72°C for 1 min), a final step of 72°C for 7 min. These set of primers quantify +e18a- and Δe18a-*Cacna1b* splice variants simultaneously. PCR products were run in 3% agarose gel stained with ethidium bromide, and densitometric analysis was performed using ImageJ [24]. This quantification method has been validated before [16]. To confirm band identity, the two bands were cloned and sequenced. To quantify total mRNA levels for *Cacna1b*, Glutamate decarboxylase-2 (*Gad-2*) and cannabinoid receptor 1 (*Cnr1*), we performed TaqMan® real-time PCR assays (ThermoFisher Scientific) with the following probes: *Cacna1b*, Mm01333678_m1; *Gad-2*, Mm00484623_m1; *Cnr1*, Mm01212171_s1; and *Gapdh*, Mm99999915_g1, which was used as constitutive control. First-strand cDNA was diluted 1:5 and 4 μL of this dilution were used in a 20 μL qPCR reaction containing Taq polymerase master mix (Applied Biosystems, 4369016) and the predesigned probes mentioned above. RT-qPCR reactions were run on an ABI 7500 Fast Real-Time PCR system (Applied Biosystems) with the following conditions: 1 cycle 95° C for 10 min, 40 cycles (95° C for 15 s and 60° C for 1 min). Each sample from at least five different mice per genotype (biological replicates) was run in triplicate (technical replicates). Ct values were determined by 7500 Software v2.3 (Applied Biosystems). Relative quantification of gene expression was performed with ΔΔ-Ct method [25].

### Statistical analysis

Two tailed unpaired Student’s t-test was performed in Excel (Microsoft).

## RESULTS AND DISCUSSION

### Validation of a mouse model to detect +e18a-*Cacna1b* mRNA in the central nervous system

To determine the localization of +e18a-*Cacna1b* mRNAs in the central nervous system (CNS), we performed a modality of *in situ* hybridization (BaseScope™). We used a mouse line with targeted deletion of e18a (Δe18a-only) to control for probe specificity. In this mouse model, DNA sequences between e18 and e19 (e18-e18a intron, e18a, and e18a-e19 intron) were removed using homologous recombination (**Fig. 1A**, [16]). To confirm deletion of e18a sequence, we performed RT-PCR in whole brain samples from WT and Δe18a-only mice. We utilized primers flanking e18a to quantify both +18a- and Δe18a-*Cacna1b* transcripts. Amplicons for each splice variant were resolved on gel electrophoresis based on size (**Fig. 1B**). In WT whole brain samples, we observed two bands, ∼290 bp and ∼230 bp, corresponding to +18a- and Δ18a-*Cacna1b* trasncripts respectively (**Fig. 1B**, *bottom left panel*, lanes 1-3). As expected, the upper band was absent in samples from Δe18a-only mice (**Fig. 1B**, *bottom left panel*, lanes 4-6). Next, we tested if the targeted deletion of e18a alters the overall *Cacna1b* mRNA levels. We quantified total *Cacna1b* mRNA using RT-qPCR with a probe that spans the splice junction between two constitutive exons, e45 and e46 (**Fig. 1B**, *top panel*). We found no significant differences in the total amount of *Cacna1b* mRNA between WT and Δe18a-only mice (Fold change relative to control ± s.e.m.: WT = 1 ± 0.12, n = 7; e18a-null = 0.89 ± 0.18, n = 7. *p* = 0.61. Two tailed, unpaired, Student’s t-test. **Fig. 1B**, *right lower panel*). Our results show that the e18a sequence was successfully eliminated from the *Cacna1b* gene and that this deletion does not alter the total *Cacna1b* mRNA levels in whole brain. Therefore, this model is ideal to control for the specificity of probes directed to e18a, thereby allowing the localization of +e18a-*Cacna1b* splice variants in CNS tissue.

### +e18a-*Cacna1b* mRNA is expressed in substantia nigra and ventral tegmental area

In the CNS, expression of *Cacna1b* is restricted to neurons, however the cell-specific expression of the *Cacna1b* splice variants has not been fully determined [4]. Previous studies using microdissections and RT-PCR showed that +e18a-*Cacna1b* mRNA is abundantly expressed in SN and VTA [17]. Furthermore, +e18a-*Cacna1b* mRNAs colocalize with tyrosine hydroxylase mRNA in both of these brain areas [17], suggesting that +e18a-*Cacna1b* mRNA is enriched in dopaminergic neurons. Here we confirmed these findings using BaseScope™with a probe designed against the e18a sequence. Briefly, ‘Z’ probes containing a short complementary region bind to the e18a sequence. This binding leads to the assembly of a signal amplification system, thereby allowing the detection of short RNA sequences (**Fig. 2A**). In our experiments, we used brain sections from Δe18a-only mice to control for probe specificity. Red signal indicates presence of +e18a-*Cacna1b* splice variants, and blue indicates counterstaining with hematoxylin (Hem) (**Fig. 2A**). Each dot detects a single mRNA molecule. We performed BaseScope™ in midbrain sections of WT and Δe18a-only mice. We compared our BaseScope™ images (**Fig. 2B**, *top panels*) to conventional in situ hybridization (ISH) images for the dopaminergic marker, *Slc6a3* (dopamine transporter, *Dat*), from SN and VTA found the Allen Mouse Brain Atlas (**Fig. 2B**, *bottom panels*) [26]. We observed that staining for e18a follows a pattern similar to *Slc6a3* in the midbrain. No signal for e18a was detected on sections from Δe18a-only mice (**Fig. 2B**, *bottom panels*). These results confirm previous studies showing that e18a is expressed in SN and VTA regions. It is important to note that we observed +e18a-*Cacna1b* expression in other cells near VTA and SNc, the identity of these cells is currently unknown. Some cells expressing +e18a-*Cacna1b* were also observed in SN pars reticulata (SNr)

**Figure 2.**
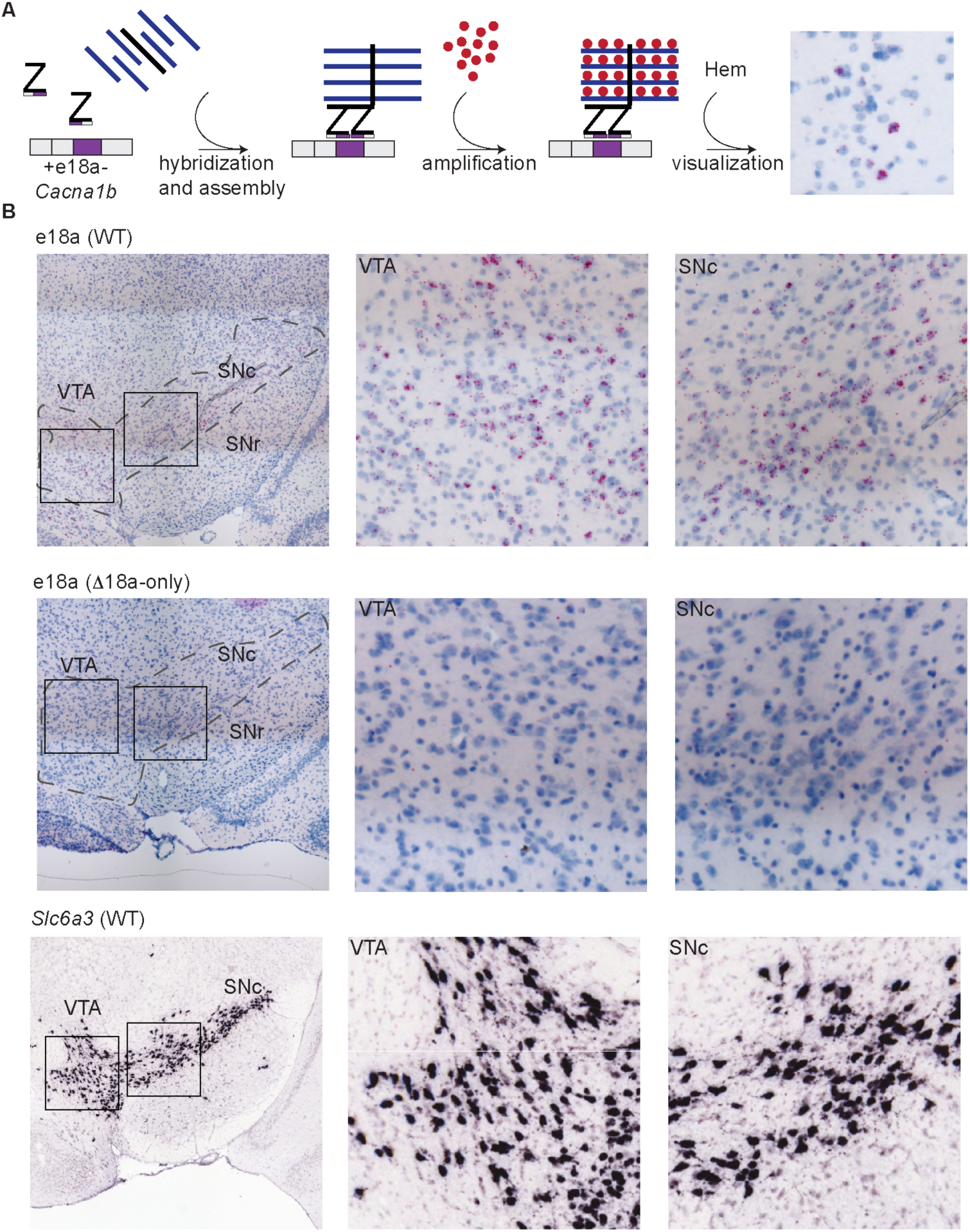
Localization of e18a-Cacna1b in dopaminergic midbrain areas using BaseScope™. **A**) Schematic of workflow for BaseScope™. Complementary Z probes were designed to target the junction e18 and e18a. These two independent Z probes bind to the *Cacna1b* mRNA in tandem, thereby allowing the assembly of an amplification complex. Subsequent signal amplification leads to the development of red coloration. Sections were counterstained with Hematoxylin (Hem). **B**) *Top panels*, BaseScope™ images of VTA and SN from WT mouse brain (*left*). Insets of VTA and SN (*middle and right, respectively*). Blue indicates all nuclei stained with Hem. Red dots indicate the presence +e18a-*Cacna1b* mRNA molecules. *Middle panels*, BaseScope™ images from areas similar to *top* panels in Δe18a-only mice. *Bottom panels*, ISH images for *Slc6a3* in SN and VTA from brain-map.org (*left*). Black squares represent sections within SN and VTA that were magnified for clarity (*middle and right*). Images credit: Allen Institute.

To our knowledge, antibodies specific for +e18a-Ca_V_2.2 channels are unavailable. This would help to convincingly show that +e18a-Ca_V_2.2 channels are present in dopaminergic neurons. However, our use of BaseScope™ with adequate negative controls shed light on the localization of e18a in the midbrain. Previous results have shown that dopamine release from VTA and SNc heavily relies on Ca_V_2.2 channels [27, 28]. Our studies and mouse models combined with previous functional studies of +e18a- and Δe18a-Ca_V_2.2 channels in mammalian systems and neurons will enable to propose further studies to unveil a potential role of e18a splicing on dopamine release.

### Expression of +e18a-*Cacna1b* mRNA in dorsal raphe nuclei and locus coeruleus

DRN and LC contain neurons that release serotonin and norepinephrine, respectively. DRN is located ventral to the cerebral aqueduct (AQ) (**Fig. 3A**, *top right panel*). Prior studies showed that +e18a-*Cacna1b* mRNA is expressed in DRN and LC [17]. We next performed BaseScope™ in these brain areas. To guide our analysis, we compared images for conventional ISH staining for the serotoninergic marker, *Slc6a4* (serotonin transporter, *Sert*), from the Allen Mouse Brain Atlas (**Fig. 3A**, *top left panel* [26]) to our BaseScope™ staining. Signal for e18a was observed ventral to AQ and followed a pattern similar to *Slc6a4*, suggesting that +e18a-*Cacna1b* splice variants are expressed in DRN (**Fig. 3A**, *middle and bottom left panels*). Sections of Δe18a-only mice in a similar area show little to no signal for e18a (**Fig. 3A**, *middle and bottom right panels*). Our results show that e18a is present in the DRN, however, further studies are needed to determine the cell-populations within the DRN that express +e18a-*Cacna1b* mRNA. Ca_V_2.2 channels are involved in the release of serotonin [27], and Ca_V_2.2-null mice show enhanced aggression that has been linked to the control of serotonin neurons excitability [29]. Our results open the possibility that splicing e18a is linked to the activity of the serotonin system.

**Figure 3.**
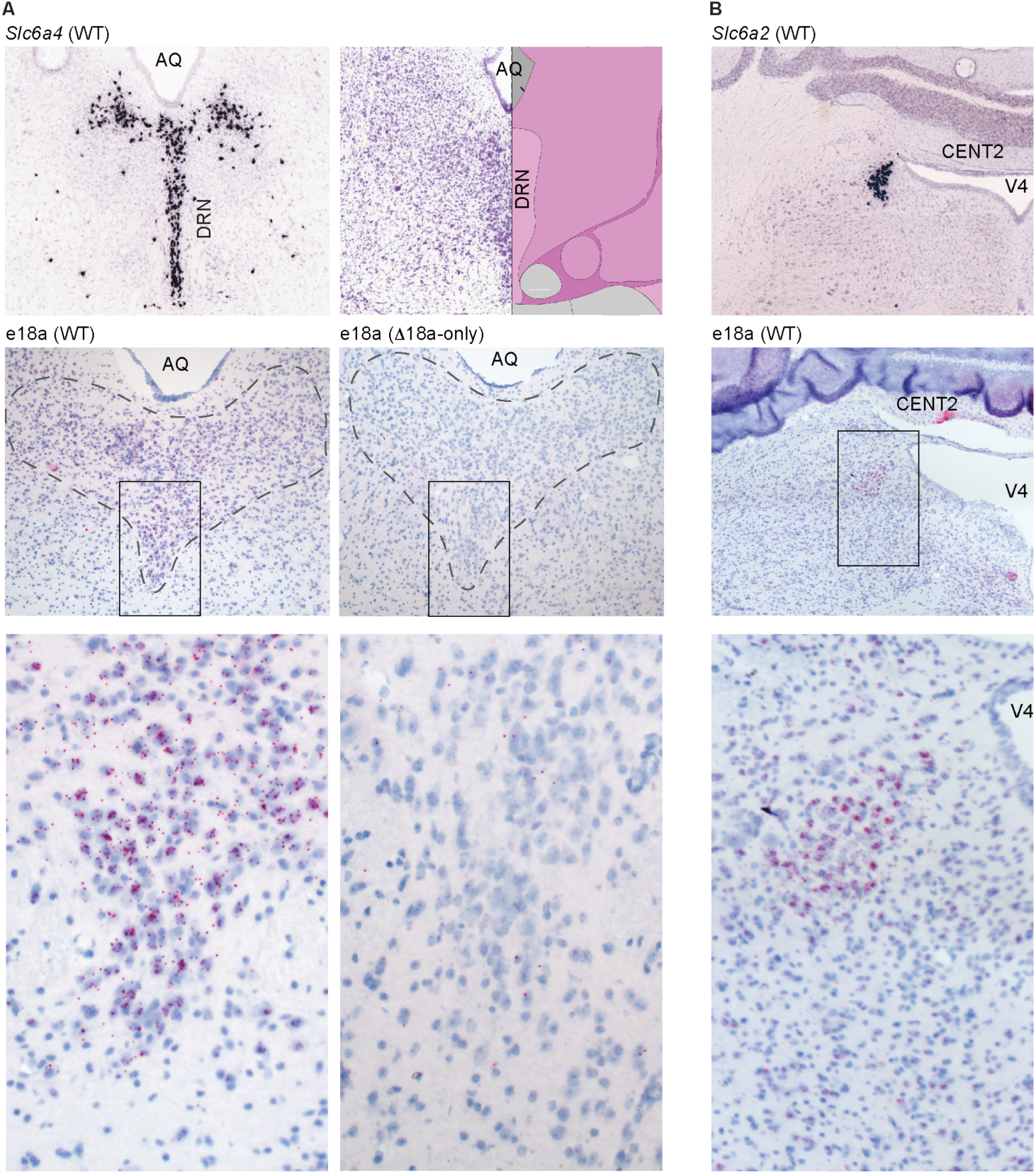
Localization of e18a-Cacna1b in the DRN and LC. **A**) *Left top panel*, ISH images for *Slc6a4* in DRN. AQ = aqueduct. *Right top panel*, Nissl staining image from brain-map.org showing the approximate location of DRN relative to AQ. Image credit: Allen Institute. *Left middle panel*, BaseScope™ images from DRN on a section of a WT mouse. *Bottom left panel*, inset shown in *left middle panel. Right middle and bottom panels*, low and high magnification (inset) BaseScope™ images of DRN sections from Δe18a-only mice. **B**) Top panel, ISH images for *Slc6a2* in LC. V4 = fourth ventricle. CENT2 = central lobule of the cerebellum 2. *Middle and bottom panels*, BaseScope™ images from WT mice in low and high magnification. Note the presence of e18a lateral to the V4. Red dots indicate the presence +e18a-*Cacna1b* mRNA. Blue denotes nuclei stained with Hem.

To determine if +e18a-*Cacna1b* mRNA is expressed in LC, we stained sections containing the fourth ventricle (V4) and the central lobule of the cerebellum II (CENT2) (**Fig. 3B**, *top panel*). We found that the signal for +e18a-*Cacna1b* mRNA is located lateral to V4 (**Fig. 3B**, *middle and bottom panels*). This signal is similar to the pattern of expression for *Slc6a2* (norepinephrine transporter, *Net*) observed in ISH of serotonin neurons images from the Allen Mouse Brain Atlas [26] (**Fig. 3B**, *top panel*). Our results suggest that e18a is present in LC, as previously reported [17].

### Distribution of +e18a-*Cacna1b* mRNA in cerebellum

In cerebellum, Ca_V_2.2 channels control the release of neurotransmitter from climbing fibers and parallel fibers synapsing onto Purkinje cells [30–32]. Ca_V_2.2 channels are also critical for the intrinsic firing of DCN neurons by coupling to calcium-dependent potassium channels [33]. In cerebellum, ∼20% of *Cacna1b* splice variants contain e18a [16]. However, the distribution of +e18a-*Cacna1b* splice variant in the cerebellum is unknown. Using the well-defined anatomy of cerebellum as landmark, we determined the expression of e18a in this area. In cerebellar cortex, little e18a signal was observed in the Purkinje cell layer (*p.c.l*.), as well as the molecular and granular layers (*m.l*. and *g.l*.) (**Fig. 4A**, *left panel, inset 1*). However, in the DCN, we observed several cell bodies stained for e18a (**Fig. 4A**, *left panel, inset 2 and 3*). Very little signal was detected in cerebellar sections from Δe18a-only mice (**Fig. 4B**, *right panel*). Our results provide a framework to test if splicing of e18a influences the firing properties of neuronal populations present in DCN.

**Figure 4.**
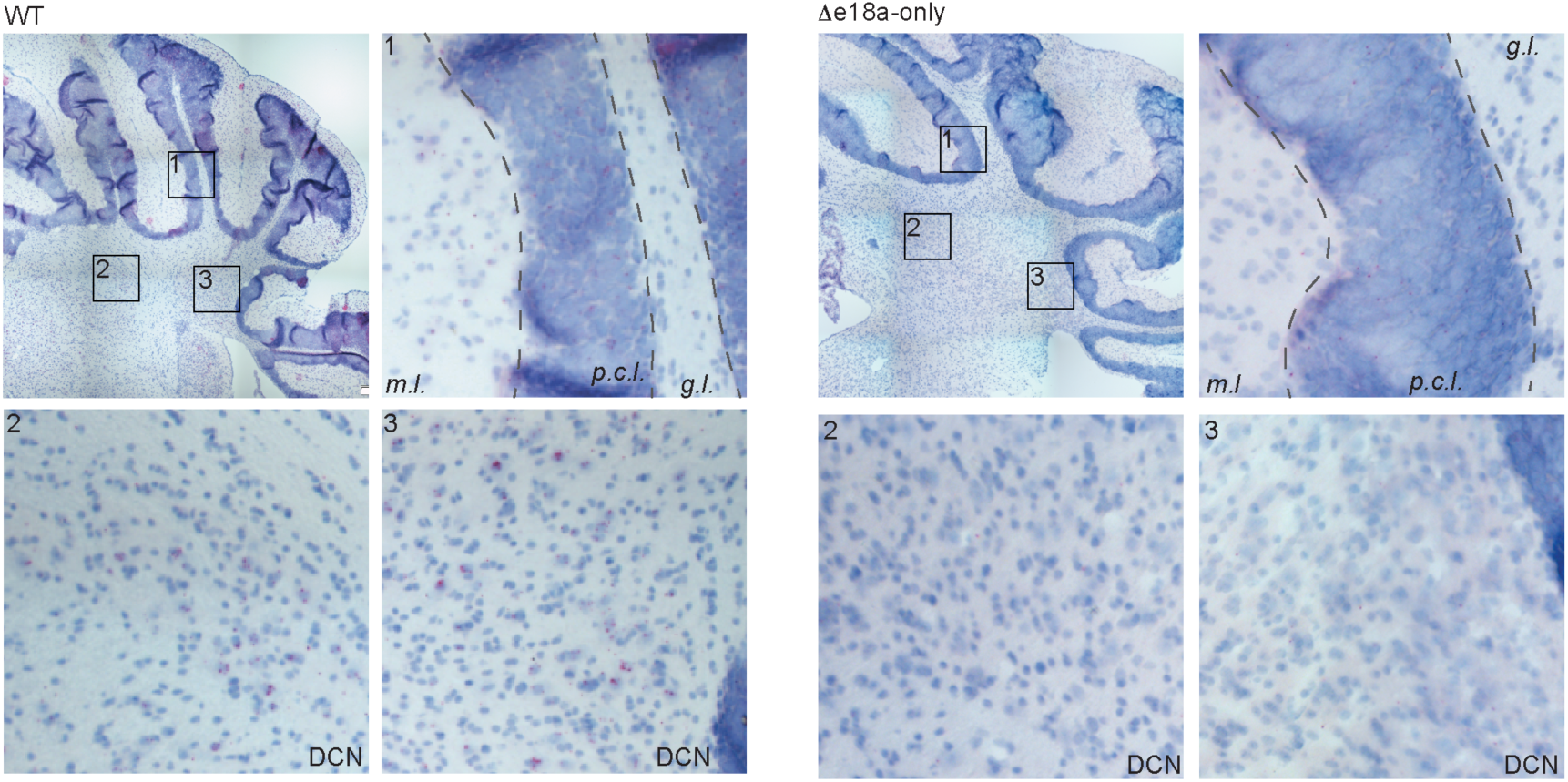
Localization of +e18a-Cacna1b in cerebellum. Left. BaseScope™ images from cerebellum of WT. Insets 1, 2 and 3 indicate magnified areas. Inset 1 represents cerebellar cortex. Layers of the cerebellar cortex are shown: molecular layer (*m.l*.), Purkinje cell layer (*p. c. l*.), and granular layer (*g. l*.). Inset 2 and 3 represent deep cerebellar areas. Red dots indicate the presence +e18a-*Cacna1b* mRNAs. Blue denotes nuclei stained with Hem. *Right*, BaseScope™ images from cerebellar areas similar to *left* from Δe18a-only mice.

### Expression of +e18a-*Cacna1b* in spinal cord

Ca_V_2.2 channels have been previously reported in spinal cord at nociceptive afferents, interneurons and motor neurons [34]. In spinal cord, we have previously shown that e18a-containing splice variants represent ∼55% of the *Cacna1b* pool of transcripts [12]. We next determined the pattern of expression of +e18a-*Cacna1b* in spinal cord. We found very little signal for e18a in laminae I-III (**Fig. 5**, *top panel, inset 1*). By contrast, large cell bodies in ventral areas (lamina IX) of spinal cord contained > 6 red dots, indicating the presence of e18a (**Fig. 5**, *top panel, inset 2*). Note the absence of e18a signal in spinal cord sections of Δe18a-only mice (**Fig. 5**, *bottom panels*). It is well known that dorsal laminae of spinal cord primarily contain neurons that receive sensory inputs from dorsal root ganglia, whereas ventral areas contain circuits that control motor neuron activity. Given that motor neurons in lamina IX have larger cell bodies relative to interneurons [35, 36], our results suggest that e18a is expressed in motor neurons. However, further studies are needed to determine if +e18a-*Cacna1b* mRNA is more abundant in motor neurons relative to interneurons of spinal cord. Previous studies have shown that some neuromuscular junctions rely on Ca_V_2.2 channels to release acetylcholine (phrenic nerve-diaphragm), whereas others utilize almost exclusively Ca_V_2.1 (sciatic nerve-tibialis muscle) [37, 38]. The cell-specific expression of +e18a-*Cacna1b* mRNA could provide an explanation for this neuromuscular junction-specific role of Ca_V_2.2 channels.

**Figure 5.**
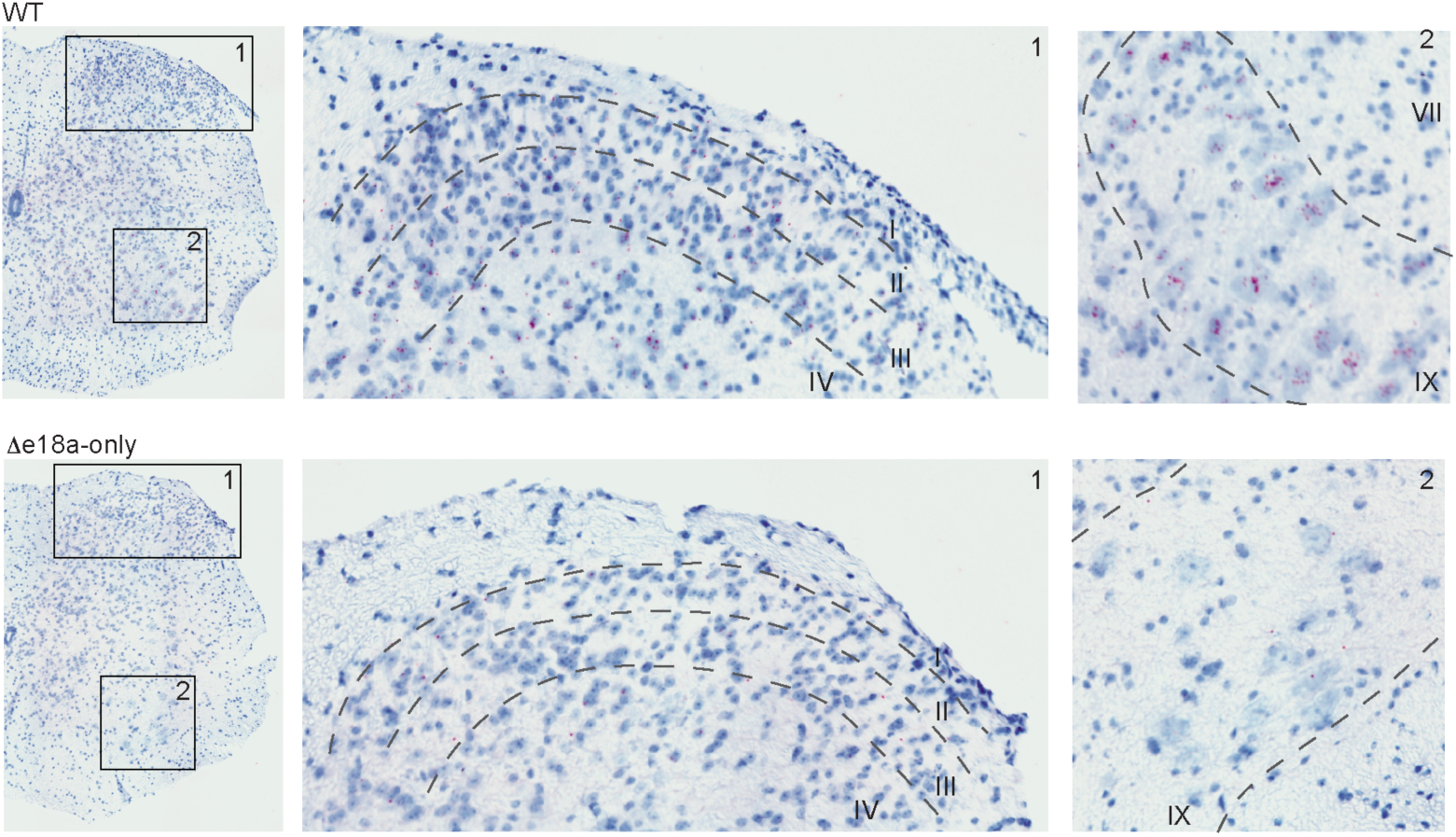
Distribution of +e18a-Cacna1b in spinal cord. Top panels, BaseScope™ images of spinal cord from WT mice. Inset 1 represents sensory areas containing laminae I-IV. Inset 2 represents motor areas containing laminae VIII and IX. Red dots are mostly localized in large nuclei in motor areas. Blue denotes counterstaining with Hem. *Bottom panels*, BaseScope™ images of similar regions as described above, but in eΔ18a-only mice.

### Expression of +e18a-*Cacna1b* splice variants in hippocampus

The functional role of Ca_V_2.2 channels in controlling neurotransmission has been extensively described in hippocampal synapses. Interestingly, the distribution of *Cacna1b* splice variants in the hippocampus is unknown. Interneurons reside in *stratum radiatum (s.r)*, whereas PNs are arranged along *stratum pyramidale (s.p.)* of both CA1 and CA3 regions. Using these anatomical landmarks, we determined the expression of e18a in CA1 and CA3 regions. Signal for +e18a-*Cacna1b* mRNA was detected in nuclei of *s.r*. and *s.p*. Interestingly, only a sub-population of *s.r*. nuclei showed strong signal for e18a in both regions (**Fig. 6A and 6B**, *left panels*). In dentate gyrus (DG), interneurons are located in the hilus (*h*.), whereas granular cells (GC) are localized in the granular cell layer (*g.c.l*). Here, we observed signal for e18a in several nuclei located in *h*. and *g.c.l*. (**Fig. 6C**). In sections from Δe18a-only mice, signal for e18a was absent throughout all regions of hippocampus analyzed (**Fig. 6A-C**, *right panels*). Based on anatomical landmarks for hippocampus, our results suggest that e18a is broadly distributed among PN, GC and a subpopulation of interneurons in the hippocampus.

**Figure 6.**
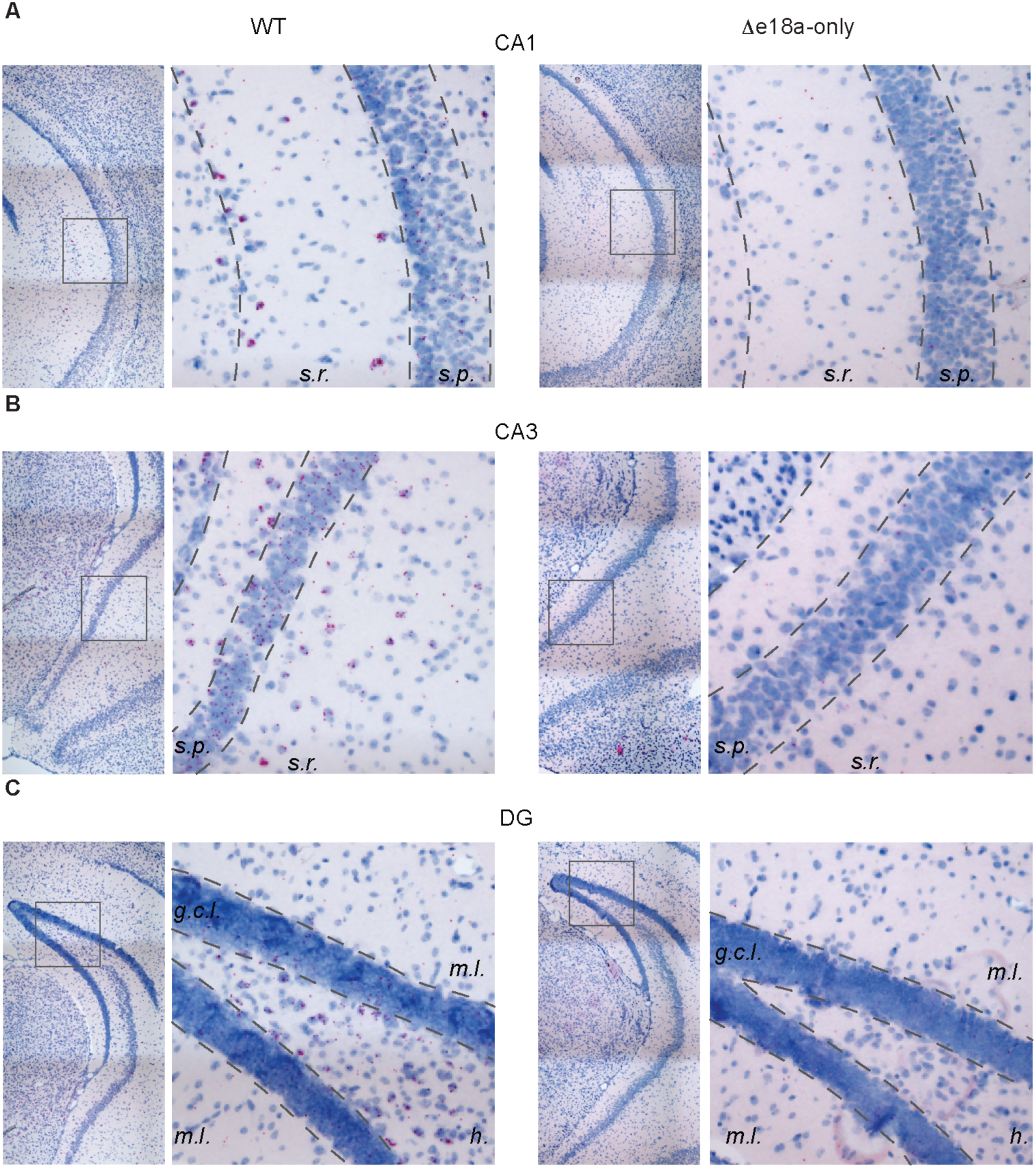
Localization of e18a-Cacna1b in hippocampus. **A**) *Left*, BaseScope™ images of CA1 region of ventral hippocampus in WT mice. Note the presence of red dots in both *stratum radiatum* (*s.r*.) and *stratum pyramidale* (*s. p*.). *Right*, similar to *left*, but in sections of Δe18a-only mice. **B**) *Left*, BaseScope™ images of CA3 region of ventral hippocampus in WT mice. *Right*, similar to *left* but in CA3 region of sections from Δe18a-only mice. **C**) *Left*, representative images of dentate gyrus of WT mice stained for e18a. DG layers are shown molecular layer (*m. l*.), hilus (*h*.), and granular cell layer (*g.c.l*.). *Right*, similar to *left* for Δe18a-only mice. In all images, red dots are indicative of the presence of e18a, and blue denotes counterstaining of nuclei with hematoxylin.

### +e18a-*Cacna1b* splice variants are enriched in cholecystokinin expressing interneurons

Ca_V_2.2 channels, together with Ca_V_2.1 and Ca_V_2.3, control transmitter release of excitatory terminals [39]. Ca_V_2.2 channels also couple to calcium-dependent potassium channels in PNs, thereby controlling neuronal firing [40]. Furthermore, CCK+INs in *s.r*. of CA1 and CA3, and *h*. of DG heavily rely on Ca_V_2.2 channels to release GABA [41]. To determine the cell-specific pattern of expression for +e18a-*Cacna1b* mRNA, we compared the relative amounts of +e18a- and Δe18a-*Cacna1b* mRNAs in CCK+INs and PNs by combining genetic labeling, FACS, and RT-PCR. To label CCK+IN, we used genetic intersectional labeling with Cre and FLPe recombinases, which resulted in the expression of tdT in CCK+INs (**Fig. 7A** and see methods). To identify PNs, mice expressing Cre recombinase from the CaMKII*α* promoter were crossed with mice containing tdT with an upstream floxed STOP codon (CaMKII*α*;tdT) [42]. Next, we performed FACS on dissociated tissue from cortex and hippocampus of CCK;Dlx5/6;tdT and CaMKII*α*;tdT mice. Total RNA was extracted, and reverse transcribed to quantify the relative amounts of +e18a- and *Δ*e18a-*Cacna1b* mRNA. We found that *Cacna1b* splice variants that contain e18a are more abundant in CCK+INs relative to CaMKII*α*^+^ PNs (% e18a relative to total *Cacna1b* mRNA, mean ± s.e.m: CCK+INs = 19.5 ± 2.5, n = 7, CaMKII*α*+PNs = 2.5 ± 1.7, n = 8. n represents the number of mice. *p* = 0.006, Student’s t-test. **Fig. 7B**). To validate the RNA extracted from sorted cell populations, we quantified *Gad-2* mRNA in both CCK+INs and CaMKII*α*+PNs. As expected, CCK+INs express higher levels of *Gad-2* mRNA relative to CaMKII*α*+PNs (% fold change, mean ± s.e.m. CCK+INs = 6.9 ± 1.57, n = 7; CaMKII*α*+PNs = 1 ± 0.41, n = 5. *p* = 0.02, Student’s t-test. **Fig. 7C**, *left panel*). Several groups have reported that CCK+INs are enriched with *Cnr1* mRNA [43–45], therefore to further validate our cell sorting, we compared the levels of *Cnr1* mRNA between CCK+INs and CaMKII*α*+PNs. We found that CCK+INs express significantly higher levels of *Cnr1* than CaMKII*α*+PNs (% fold change, mean ± S.E. CCK+IN = 102 ± 23.0, n = 7, CaMKII*α*+PNs = 1.0 ± 0.30, n = 8, p = 0.0002, Student’s t-test. **Fig. 7C**, *right panel*). Taken together our results strongly suggest that +e18a-*Cacna1b* pre-mRNA is expressed at higher levels in CCK+INs relative to CaMKII*α*+PNs.

**Figure 7.**
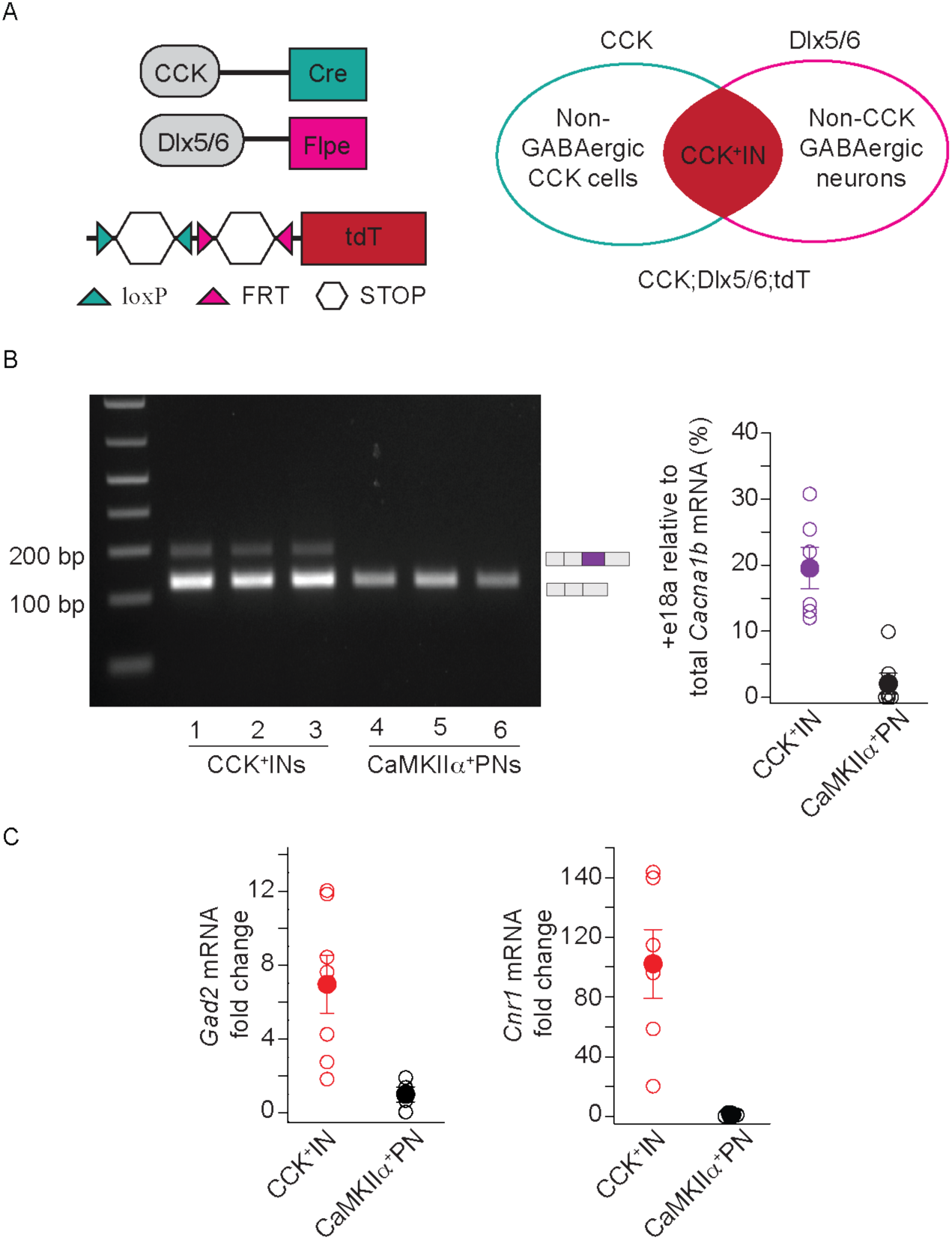
+e18a-Cacna1b mRNA is enriched in CCK+INs. **A**) Intersectional genetic labeling of CCK+INs. Cre recombinase is expressed from the CCK promoter active in CCK interneurons and projection neurons, whereas Flpe recombinase is expressed from the Dlx5/6 promoter active in forebrain GABAergic interneurons. The tdT allele present in Ai65-D mouse contains two STOP codons (white hexagons) flanked by loxP and FRT sites. In a mouse with these three alleles, the combined action of Cre-loxP and Flpe-FRT systems removes both STOP codons allowing expression of tdT in GABAergic CCK+INs, but not in PNs. **B**) *Left panel*, representative gel of RT-PCR in RNA purified from sorted CCK+INs and CaMKII*α*+PNs. *Right panel*, comparison of +e18a-*Cacna1b* relative to total *Cacna1b* mRNA in CCK+IN and CaMKII*α*+PN. **C**) Validation of sorted cells. Total amounts for *Gad2* and *Cnr1* mRNA were compared between CCK+INs and CaMKII*α*+PNs using RT-qPCR. Data are shown as mean (filled symbols) ± s.e.m., and the averages of individual biological replicates (empty symbols).

CCK+INs are linked to mood disorders [46, 47], express relatively high levels of the cannabinoid receptor protein (CB1R) [48], and contribute to the behavioral tolerance of tetrahydrocannabinol [49]. Given that CB1R agonists downregulate Ca_V_2.2 channels to inhibit GABA release in CCK+INs, our findings suggest an anatomical link between e18a splicing in *Cacna1b* and the effects of CB1R agonists of transmitter release. Furthermore, CCK+INs in hippocampus show robust asynchronous release and spontaneous release that relies on Ca_V_2.2 channels [41]. Due to the slower inactivation rates and positive shift in voltage-dependent inactivation of +18a-Ca_V_2.2 channels relative to Δ18a-Ca_V_2.2 channels, it is possible that inclusion of e18a in this neuronal type enhances asynchronous and spontaneous release.

## ABBREVIATIONS

VTA, SN, DRN, LC, DAT, DCN, CA1, CA3, SNc, GCL, PN, CCK, MCL, PCL, CB1R, CNR1, GAD2, SR, SP, H, GAPDH.

## ACKNOWLEDGEMENTS

This work was supported by the National Institute of Mental Health [grant number, R00MH099405]. We thank Sylvia Denome for generating the Δ18a-only mice, Diane Lipscombe lab for their advice on BaseScope™, and Shayna Mallat for meaningful comments on this manuscript.

